# STRING-ing together protein complexes: corpus and methods for extracting physical protein interactions from the biomedical literature

**DOI:** 10.1101/2023.12.10.570999

**Authors:** Farrokh Mehryary, Katerina Nastou, Tomoko Ohta, Lars Juhl Jensen, Sampo Pyysalo

**Author notes:** Equal contribution (Farrokh Mehryary, Katerina Nastou).

## Abstract

Understanding biological processes relies heavily on curated knowledge of physical interactions between proteins. Yet, a notable gap remains between the information stored in databases of curated knowledge and the plethora of interactions documented in the scientific literature. To bridge this gap, we introduce ComplexTome, a manually annotated corpus designed to facilitate the development of text-mining methods for the extraction of complex formation relationships among biomedical entities. This corpus comprises 1,287 documents with ∼3, 500 relationships. We train a novel relation extraction model on this corpus and find that it can highly reliably identify physical protein interactions (F1-score=82.8%). We additionally enhance the model’s capabilities through unsupervised trigger word detection and apply it to extract relations and trigger words for these relations from all open publications in the domain literature. This information has been fully integrated into the latest version of the STRING database, and all introduced resources are openly accessible via Zenodo and GitHub.

## Introduction

The study of physical protein interactions forms a basis for understanding biological processes. These interactions are captured from experimental data [Oughtred et al., 2021, Orchard et al., 2014, Licata et al., 2012] or scientific articles [Meldal et al., 2019, Giurgiu et al., 2019, Gillespie et al., 2022] and are provided as periodically curated databases.

These curation efforts have been complemented by co-occurrence-based text mining in protein interaction databases, such as STRING [Szklarczyk et al., 2023] and HumanNet [Kim et al., 2021], to obtain more comprehensive networks. While this approach is powerful for linking molecules that function together, the fact that proteins are mentioned together in text is not enough to infer that they also physically interact. Thus, earlier versions of STRING [Franceschini et al., 2012] employed a rule-based system to help with the extraction of such interactions.

In the field of biomedical natural language processing (BioNLP), the past two decades have witnessed significant progress, driven by the development of more sophisticated and accurate deep learning-based methodologies [Milošević and Thielemann, 2023] and manually annotated corpora. Training deep learning models typically involves a two-step process: self-supervised pre-training on an unannotated corpus with a language modeling objective, and fine-tuning on manually annotated data for a specific downstream task (e.g. relation extraction). Models based on the transformer architecture [Vaswani et al., 2017] such as BERT [Devlin et al., 2019], have been particularly effective, combining efficient training of large-scale models using GPU acceleration with state-of-the-art performance across a broad range of tasks.

The effectiveness of the original BERT and other biomedical domain-specific BERT models [Lee et al., 2019, Lewis et al., 2020] depends on the availability of manually annotated corpora for fine-tuning. Generating these corpora requires expert knowledge, making it a costly endeavor. Particularly in the domain of physical protein interactions, existing manually annotated corpora are typically tailored to a specific task [Pyysalo et al., 2008], and the differences in the definition of complex formation, among these corpora, pose a substantial challenge in interoperability. These differences underscore the necessity of an annotated corpus tailored explicitly to STRING database needs, as leveraging existing data proves challenging given the varied definitions.

In this study, our primary objective was to develop a system to support the relation extraction of physical `protein-protein` interactions from the literature for the STRING database. For this purpose, we present ComplexTome, a new corpus annotated with complex formation relationships between biomedical entities. We have also built a relation extraction pipeline, trained it on ComplexTome to extract such relationships from the openly accessible biomedical literature, and created a trigger word detection system to aid in interpreting the results. We provide the corpus, code, and all results produced by the large-scale runs of our system via Zenodo https://doi.org/10.5281/zenodo.10693924, Github https://github.com/farmeh/ComplexTome_extraction, and the latest version of STRING https://string-db.org/.

## Materials and methods

### The ComplexTome corpus

#### Document selection for corpus annotation

The selection of documents for ComplexTome involved a three-step approach of selecting documents from:

1. **Existing corpora**: Initially, we explored established corpora, particularly the BioNLP ST 2009 training and development datasets [Kim et al., 2009]. From these datasets, we identified and selected a total of 135 abstracts featuring instances of complex formation events, from which we could potentially extract documents with the desired relationship type present. As the definition and annotation of binding events for the BioNLP ST 2009 corpus were not compatible with the relationship annotation we were aiming for in ComplexTome, all existing annotations were discarded and the documents selected were reannotated from scratch.
2. **Resources enriched in positive relationships**: To broaden the corpus, we curated 400 abstracts extracted from a collection of 66,757 publications used as evidence to support physical or genetic interaction entries in the BioGRID [Oughtred et al., 2021], IntAct [Orchard et al., 2014] and MINT [Licata et al., 2012] interaction databases. Additionally, 400 paragraphs extracted from 12,577 articles available as PubMed Central Open Access (PMC-OA) full-text articles were selected from the same databases. Documents used to annotate more than 20 interactions in the databases were removed from the selection pool.
3. **Resources enriched in negative relationships**: We also selected 300 abstracts extracted from 21,941 papers used for pathway annotation in the Reactome pathway knowledgebase [Gillespie et al., 2022] and 50 abstracts extracted from 15,319 papers in BioGRID filtered to include only experimental associations for genetic interactions.

During steps 2 and 3 we used a dictionary-based Named Entity Recognition (NER) method [Jensen, 2016], to detect protein entities within the large document pools and restricted the selection to abstracts containing a moderate number of detected protein entities (between 2 and 40). To prevent over-representation of commonly mentioned protein entities, abstracts featuring highly mentioned proteins were limited to comprise at most 2% of the selected documents. All documents in ComplexTome were annotated using the BRAT rapid annotation tool [Stenetorp et al., 2012].

#### Named Entity Annotation

We annotated four Named Entity (NE) types, namely `Gene or Gene Product (Protein` hereafter), `Chemical` which encompasses standalone chemicals that are not part of larger chemical/protein entities, `Complex` for entities describing stable assemblies of two or more macromolecules, in which at least one component is a protein, and `Protein_Family`, for entities which represent an evolutionarily conserved group of gene/proteins or a group of entities with the same function. To assist the manual annotation process of NEs, we employed automated NER [Jensen, 2016] for the detection of Protein entities in our corpus.

In the scientific literature, it is common to encounter alternative names referring to identical entities. We have systematically recorded these equivalent names in ComplexTome. This practice is crucial for accurate evaluation, as it allows relationships stemming from either entity to be recognized as valid [Kim et al., 2009]. To allow for easy filtering of the NEs, we used five entity attributes to mark NEs in our corpus: “Mutant”, “Fusion”, “Non-coding”, “Small protein post-translation modification” and “Blocklisted”. For a description of these attributes please refer to our annotation documentation https://katnastou.github.io/annodoc-physical-protein-interaction-corpus/.

#### Relationship annotation

In ComplexTome, we identified explicit mentions of physical protein interactions and annotated these as undirected binary relationships with the type `Complex_formation`. We added annotations for any statement implying the existence of a complex, but not statements explicitly denying the formation of a complex. A relationship annotation example is shown in Figure 1.

**Fig. 1.**
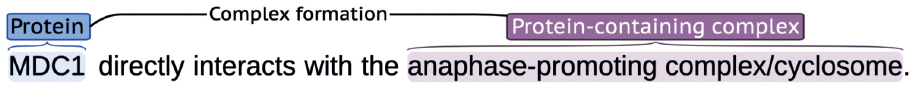
Illustration of the relationship representation in ComplexTome. A Complex formation relationship between a Protein (”MDC1”) and a Complex (”anaphase-promoting complex/cyclosome”) participant has been identified by the annotators in the example above.

The annotations were performed by two domain experts, ensuring accurate relationship identification. Moreover, to establish consistent annotation guidelines and maintain annotation quality, we conducted an Inter-Annotator Agreement (IAA) analysis by independently annotating a set of abstracts during the two initial steps of document selection. The process encompassed four rounds of independent annotations with 20 documents selected and annotated per round. We calculated the F1-score of IAA after each round to assess annotation consistency and corpus quality. For comprehensive details on the annotation rules followed to produce the corpus, we refer readers to the annotation documentation provided to the annotators (See Section 2.1.2). This documentation served as a reference to ensure a shared understanding of the rules among annotators and contributed to maintaining the overall quality of annotations.

### Relation extraction system

We have developed a relation extraction system that is capable of extracting `Complex_formation` relations between `Protein` named entities, as stated in biomedical texts. We cast the task of relation extraction as binary classification, predicting whether a `Complex_formation` relation has been stated for the two candidate NEs in the given input text (i.e. a positive label), or not (i.e. a negative label).

The system is based on deep neural networks with an architecture consisting of a pre-trained transformer encoder, followed by a decision layer with a softmax activation function. The system can utilize existing pre-trained language models that are currently available in the Hugging Face repository (https://huggingface.co/), allowing us to benchmark different models and use the best available model for the task. Training, validation and prediction files can be provided to the system in BRAT standoff format (as well as a custom JSON format). The system supports two input representation methods: marking and masking the NEs [Mehryary et al., 2020] and it can be trained with a wide variety of hyper-parameters including maximum sequence length (`MSL`), learning rate, mini-batch size, number of training epochs and random seed. During the training on the ComplexTome training data, pre-trained weights of the encoder are fine-tuned, while randomly initialized weights (such as the weights of the decision layer) are learned from scratch. After each training epoch, evaluation metrics are calculated on the development set and used for hyper-parameter optimization. We do not use any early stopping rule, instead, we train a network for the specified number of epochs (given as a hyper-parameter) but use model weights of the epoch which has yielded the highest F1-score. For implementation, we use Python programming language with `TensorFlow` and `transformers` libraries.

#### Preprocessing, input representation and example generation

As biomedical texts typically contain more than one candidate NE pair and can be very long, not fitting within the maximum input tokens of transformer models, we process each input document as follows.

1. **Marking or masking the entities**: To inform the classifier which two particular NEs constitute a pair at-a-time for label prediction, we transform the text by encoding the entities in the input document, either using a marking approach or a masking approach, utilizing the language model’s “unused” tokens for this aim. Additionally, we prepend a [`CLS`] token, representing the snippet start, and append a [`SEP`] token to the tokens, representing the end of snippet, before feeding them to neural network models.
2. **Tokenization, distance calculation, and example generation**: For each candidate NE pair, after transforming the text (marking/masking), we tokenize the text, and if the two entities can fit into a window with a size less than or equal to the specified `MSL`, we generate a machine learning example for the pair. The pair can have a positive, or a negative label (for training) or can be unlabeled (during prediction).

Our pre-processing approach provides two benefits: since we are not performing any sentence boundary detection, the system is able to train with and predict cross-sentence relations. In addition, there will be no problem in dealing with long texts, since we rely on a window (sequence of sub-tokens) that can always be fed to the transformer encoder. Supplementary Material Section 1 provides more details and a few examples for all steps described here.

#### Experimental setup

We performed a document-based split of ComplexTome to generate separate training, development, and test sets, which consisted of 772, 258, and 257 documents respectively. We train the system on the training set and optimize it on the development set. We use a grid search to try different transformer models and find the optimal values of hyper-parameters. To minimize the effect of initial random weights on evaluation scores [Mehryary et al., 2016], we repeat each experiment four times and compare different experiments based on the average and standard deviation of the F1-scores. Each *experiment* consists of training a relation extraction system with different initial random weights but with the same transformer encoder weights and the exact set of hyper-parameters on the training set and evaluating the model on the development set. The held-out test set was only used once for the final evaluation of our best system. Supplementary Material Section 2 provides a list of tested models and hyper-parameters.

Even though our corpus contains four NE types (`Protein, Chemical, Complex` and `Protein_Family`), and `Complex_formation` relationships can occur between any two NEs, for the real-world application for which the system was intended for, i.e. extracting `Protein-Protein` interactions for STRING database v12, the system had to deal with texts including only `Protein` entities, which are not “blocklisted”. Hence, to have a realistic optimization approach for large-scale prediction, we filtered out all `non-Protein` entities and `Protein` entities with the “blocklisted” attribute (and their relations) from the development and test sets. By contrast, in order to assess whether we could leverage the full potential of relationship annotation in our corpus, during training we performed several experiments with different training schemes. More details on the experimental setup are provided in Supplementary Material Section 3.

### Trigger detection system

While detecting `Complex_formation` relations between biomedical entities in the scientific literature is a task of paramount importance to the scientific community, it is even more beneficial if, for each `Complex_formation` relation extracted from a text snippet, there was a system that could also highlight the most important word or phrase in the text snippet that explicitly or implicitly denotes the relation. In the BioNLP community, such words or phrases are called “trigger words” (hereafter “triggers”) and were popularized as part of the biomedical event representation of the BioNLP Shared Task 2009 on Event Extraction [Kim et al., 2009]. In the context of `Complex_formation` extraction, triggers can be as simple as “bind", or as complicated as “tandem affinity purification".

Traditionally, “trigger detection” (automatic recognition of the keywords/phrases behind the extraction of events or relations) has been defined as a supervised NER task, relying on manually annotated data for training and evaluation. However, with the help of model explanation methods, such as *Integrated Gradients* [Sundararajan et al., 2017] and *SHAP (SHapley Additive exPlanations) values* [Lundberg and Lee, 2017], we aim to automatically find the triggers in an unsupervised fashion. The general idea here is to apply such methods to calculate and assign a score to each token of the input with regard to its contribution to the predicted label (`Complex_formation` in our case), and by ranking the tokens based on their scores, we can obtain the token(s) with the highest contribution to the label for the extracted relations. We hypothesize that these tokens will frequently correspond to the triggers. Naturally, this is done only for `Protein-Protein` pairs with a positive label, i.e. when the model has predicted that there is a `Complex_formation` relationship between two candidate NEs in a given input text, as by definition, triggers are the words or phrases that explicitly state or imply the existence of a relationship between two NEs.

While using model explanation methods to attribute importance scores to input tokens has previously been applied in areas like sentiment analysis (e.g. identifying keywords in movie reviews[Dewi et al., 2022]), its application to automatic trigger detection, as presented in this manuscript, is novel.

#### Experimental setup

Model explanation methods are generally used to provide useful insights about how a *trained* model works. Therefore, the design and development of the trigger detection system started once we had obtained the final neural network model for `Complex_formation` extraction.

While such methods are unsupervised, not requiring any manually annotated data for training, we still needed manually annotated data to evaluate and compare approaches, since there is no guarantee that the token(s) with the highest contribution to a positive label is the trigger we aim to recognize. Therefore, we focused on the **positive pairs** of the ComplexTome development set. We first split this set into two equally sized sets (based on the number of documents), hereafter a **trigger development set**, and a **trigger test set**. We then excluded those infrequent positive examples from the two sets that do not fit into a window of 128 tokens, since this is the best MSL found during hyper-parameter optimization, and our best model has been trained with this restriction for real-world application. After filtering, there are 284 and 275 positive `Protein-Protein` pairs in the trigger development and test set respectively, for which we aim to recognize triggers.

The two sets were subsequently given for annotation to an expert, with the annotation task of highlighting trigger text spans for each positive pair. If two or more text spans were considered as equivalently valid triggers, they were all annotated as triggers for a `Protein-Protein` pair, but we emphasize that correctly recognizing and showing only one of the triggers to the end-user in such cases is sufficient. The section “Trigger word annotation” in the annotation guidelines (https://katnastou.github.io/annodoc-physical-protein-interaction-corpus/) provides details with interesting examples.

#### Trigger detection methods

We experiment with two commonly used model explanation methods:

1. **Layer Integrated Gradients (**`LIG`), as implemented in the Captum library (https://captum.ai/), which is a variant of the Integrated Gradients method that assigns an importance score to a desired layer’s outputs of a trained neural network model, for every token in the input snippet.
2. **SHapley Additive exPlanations (**`SHAP`**) values** (https://github.com/shap/shap), which is a method commonly used for explaining the predictions of a machine learning model. `SHAP` calculates a value that represents the contribution of each token in the input snippet to the model outcome, thus the values reflect the importance of each token with regard to the label.

Our best relation extraction model was obtained by fine-tuning a pre-trained RoBERTa model [Lewis et al., 2020] on the ComplexTome training set. This is consistent with the top-performing teams in the recent DrugProt relation-extraction track of BioCreative VII using pre-trained RoBERTa models [Miranda-Escalada et al., 2023]. By feeding trigger development set examples as input to the model and applying the LIG method on the outputs of the embedding layer and the 24 hidden RoBERTa layers in this model, we obtain 25 vectors for each development set example. By discarding [CLS], [SEP], and “unused” tokens and then simply choosing the token with the highest score in each vector as the predicted trigger, we obtain 25 different predictions for each development set example. Therefore, we form 25 different prediction sets of the development set (each based on a particular layer in the model), which can further be evaluated against the gold standard. Similarly, by applying the `SHAP` method, we obtain one set of predictions. Initial experiments showed slight differences when feeding the inputs *with* and *without* the [CLS] and [SEP] tokens. Therefore, we tried both types of inputs for the `SHAP` method. In total, we obtained 27 predicted sets for the development set triggers, which we then compared against the gold standard and calculated evaluation scores. To further improve the results, we also defined a set of post-processing heuristic rules, targeting and removing irrelevant tokens from the vectors before choosing the tokens with the highest score as predicted triggers. Supplementary Material Section 4 provides further details about our trigger detection methods.

### Baseline method

For comparison, we also develop a simple baseline method. In this method, we first obtain the list of all words or phrases that are highlighted as triggers in the trigger development set. Then, for each `Complex_formation` relation in the trigger development set, we define two windows around the two candidate NEs (the window size in sub-tokens is an optimizable parameter, ranging from 1 to the maximum possible value for the trigger development set). Based on the selected window size, for each word or phrase in the list, we search the windows with regular expressions, and if we can match a word or phrase against a span in the texts of two windows, we annotate the span as a recognized trigger.

## Results and discussion

### Corpus statistics

ComplexTome is a high-quality corpus — supported by the fact that we attained over 90% agreement over four rounds of IAA — comprising 1,287 documents with approximately 300,000 words. There are 3,486 `Complex_formation` relationships in the corpus, ensuring a rich and diverse collection of relations for training neural network models for the relation extraction task. Over 96% of these relationships are intra-sentence (i.e. within sentence boundaries), while less than 4% cross sentence boundaries. There is a large number of NEs in the corpus, namely 20,228 `Protein, 2,185 Chemical, 1,500 Complex`, and 3,019 `Protein_Family` entities. Notably, we gave the “Blocklisted” attribute to 795 entities during annotation and later properly filtered them out for the training and development of the relation extraction model.

### Relation extraction

#### System evaluation

We used grid search to find the optimal values of hyper-parameters and compare different pre-trained transformer encoders, using the methodology described above. Our best result was achieved using the `RoBERTa-large-PM-M3-Voc` encoder [Lewis et al., 2020], a 24 layer RoBERTa-based language model, pre-trained on biomedical and clinical texts, and using the following hyper-parameter values: *MSL* = 128, *lr* = 3*e −* 6, *training epochs* = 11, *minibatch size* = 5. This experiment resulted in 86.75% average precision, 82.05% average recall, and 84.28% average F1-score on the ComplexTome development set. Table 1 shows the details of the four models used in this experiment and the evaluation scores measured on the development set.

**Table 1.**
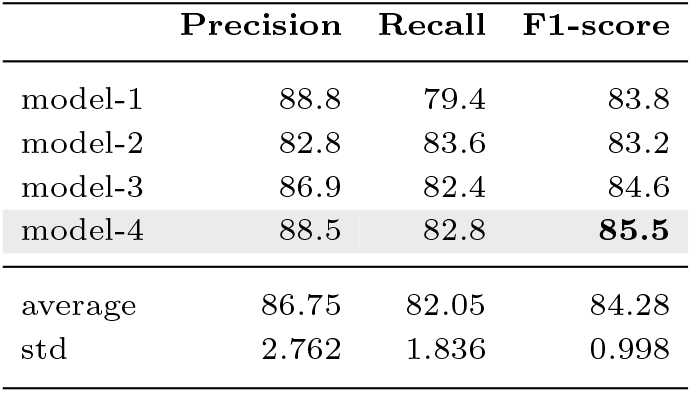
Performance of the best experiment on the ComplexTome development set. The best model (highlighted in gray) is used for large-scale prediction on the entire literature.

|  | Precision | Recall | F1-score |
| --- | --- | --- | --- |
| model-1 | 88.8 | 79.4 | 83.8 |
| model-2 | 82.8 | 83.6 | 83.2 |
| model-3 | 86.9 | 82.4 | 84.6 |
| model-4 | 88.5 | 82.8 | **85.5** |
| average | 86.75 | 82.05 | 84.28 |
| std | 2.762 | 1.836 | 0.998 |

Our best model achieves 85.5% F1-score on the development set and 87.3% precision, 78.8% recall and 82.8% F1-score on the held-out test set. We have selected this particular model for large-scale execution and for providing the text-mined associations for the physical subnetwork of STRING v12.

#### Manual error analysis

We present an analysis of the errors produced by the best relation extraction model on the test set, grouped into categories in Table 2. For a detailed overview of all errors, please refer to Supplementary Material Section 5.

**Table 2.** Error analysis for the relation extraction system.

| Error Type | Count |  |  |
| --- | --- | --- | --- |
|  | FP | FN | Total |
| Ambiguous keyword | 18 | 26 | 44 |
| Rare keyword | 2 | 23 | 25 |
| Convolved text excerpt | 15 | 39 | 54 |
| Co-reference resolution | 11 | 9 | 20 |
| Annotation error | 11 | 8 | 19 |
| Total | 57 | 105 | 162 |

\*FN: false negative, FP: false positive

We identified five error categories, with none appearing to be the primary cause of errors observed in the test set. The first category is “ambiguous keyword", which involves words like “association” that can describe both physical interactions and other types of relationships. This makes it hard to assign an accurate relationship label, resulting in both FPs and FNs. “Rare keyword” pertains to relationships between NEs that annotators have identified based on their biological knowledge, but which are indicated by phrases or words that are seldom encountered in the literature (e.g. non-covalent association) and thus result mostly in FNs. “Convoluted text excerpt” refers to text segments characterized by syntactic intricacies, including complex sentences and cross-sentence relationships, which are inherently difficult to understand, in some cases even for humans. A separate error type related to this is “co-reference resolution". These errors arise when it is unclear, based on syntax, which subject(s) a specific `Complex_formation` relationship corresponds to, and produce FPs as well as FNs.

Finally, “annotation error” corresponds to cases where the annotators have made specific mistakes — often stemming from ambiguity — and these require fixing upon clearer examination. Recalculation of the statistics after excluding annotation errors lead to an increase in precision (89.7%), recall (80.1%) and F1-score (84.7%), representing a slight improvement compared to the original observations on the test set.

### Trigger detection

#### System evaluation

For large-scale relation extraction, we had trained a TensorFlow-based model that achieved 88.5% precision, 82.8% recall, and 85.5% F1-score on the ComplexTome development set. Since the Captum implementation of the LIG method operates only on models that are trained with the PyTorch library, we trained another relation extraction system (with the same `RoBERTa-large-PM-M3-Voc` encoder and the same best hyper-parameters) using the PyTorch library and during the implementation of the code, we also fixed a couple of minor implementation errors. This resulted in a better relation extraction model with 88.8% precision, 85.4% recall and 87.1% F1-score on the development set. We used this model for developing and optimizing our trigger detection system, and also for large-scale execution of the trigger detection system on biomedical literature.

Trigger detection methods are usually evaluated with the standard metrics used for NER tasks (precision, recall, and F1-score). However, annotation of the trigger development set showed that: (1) a trigger can span multiple input tokens, and (2) there can be more than one alternative trigger spans that are equally valid to annotate for a `Complex_formation` relationship (for more details please refer to our annotation guidelines). However, from the practical application standpoint, it is good enough that we recognize only a part of a multi-token trigger. Similarly, if there are alternative trigger spans for the same `Complex_formation` relationship (e.g. The CD40*-*TRAF2 *interaction*), recognizing one of them is sufficient. Therefore, as evaluation metrics for method development and optimization, we take average of precision scores, average of recall scores, and average of F1 scores for left-bound, right-bound, overlap and exact matching of detected trigger spans against the gold-standard annotations. These measures penalize the method when it fails to detect alternative trigger spans for a single `Complex_formation` relationship, but since they were easy to program and calculate, we used them for method development. For the final evaluation of our best method, we used manual evaluation, not penalizing for such cases.

For the development and optimization of the trigger detection system, we evaluated our methods on the positive `Protein-Protein` pairs in trigger development set. An initial experiment with the `LIG` and `SHAP`-based methods showed that when the relation extraction model makes a mistake and produces a negative label for a positive pair (i.e. the FN predictions of the relation extraction system), any effort in detecting the trigger by model explanation methods leads to mistakes, producing incorrect triggers (FP trigger spans). For example, the `SHAP` method (with [CLS] and [SEP] tokens, and without the post-processing heuristics) results in 62.08% overlap precision, 55.45% overlap recall, 58.58% overlap F1-score, but if we first discard all FN pairs, then the same method results in 67.69% precision, 52.73% recall and 59.28% F1-score. From the perspective of end-users, having a higher precision is very important, because it will increase the credibility of text mining results. Therefore, for `LIG` and `SHAP`-based methods, we chose to always check the relation label as predicted by the trained RE model, and only pursue trigger detection if a positive label has been predicted by the model. Table 3 shows the results of our best approaches on the trigger development set, before and after applying post-processing heuristics.

**Table 3.** Comparison of trigger detection methods on the trigger development set. The best method (highlighted in gray) is used for large-scale execution on the entire literatur/e=

|  | overlap<br>Pre | overlap<br>Rec | overlap<br>F1 | exact<br>Pre | exact<br>Rec | exact<br>F1 | avg<br>Pre | avg<br>Rec | avg<br>F1 |
| --- | --- | --- | --- | --- | --- | --- | --- | --- | --- |
| LIG (best layer=9) | 58.36 | 49.70 | 53.68 | 53.02 | 45.15 | 48.77 | 55.69 | 47.43 | 51.23 |
| Baseline | 50.12 | 65.15 | 56.65 | 48.25 | 62.73 | 54.55 | 49.18 | 63.94 | 55.60 |
| SHAP (with [CLS] and [SEP]) | 62.21 | 55.76 | 58.81 | 56.86 | 51.52 | 54.05 | 59.53 | 53.79 | 56.52 |
| SHAP (without [CLS] and [SEP]) | 62.71 | 56.06 | 59.20 | 58.64 | 52.42 | 55.36 | 60.68 | 54.24 | 57.28 |
| SHAP + Heuristics (without [CLS] and [SEP]) | 83.13 | 62.42 | 71.31 | 73.90 | 55.76 | 63.56 | 78.41 | 59.09 | 67.40 |
| SHAP + Heuristics (with [CLS] and [SEP]) | 85.20 | 63.64 | 72.86 | 74.80 | 56.67 | 64.48 | 79.90 | 60.31 | 68.73 |
| LIG (best layer=14) + Heuristics | 95.12 | 70.91 | 81.25 | 85.77 | 63.94 | 73.26 | 90.45 | 67.43 | 77.26 |

\*Pre: Precision, Rec: Recall, F1: F1-score, avg: average of left-bound, right-bound, overlap and exact matching scores

As Table 3 shows, the baseline method performs very poorly, achieving 55.60% average F1-score, showing that a simple pattern matching approach is not good for trigger detection. This is mostly due to the high amount of FP spans that resulted in the lowest average precision among all methods.

Another interesting observation is that the `LIG` method (without post-processing heuristics) performs the worst. Although having a higher precision than the baseline, this method yields a much lower recall, and lower F1-score, suggesting that the LIG method without additional heuristics is not fit for the job. In contrast, the two SHAP-based methods (without the heuristics) have performed closely, with 56.52% and 57.28% F1-score, and slightly outperformed the baseline, showing that “right out of the box” and without additional tweaking, vanilla `SHAP` methods outperform the `LIG` method on the task.

The post-processing heuristic rules significantly improve the results, increasing both precision and recall, which is evident in both `SHAP` and `LIG`-based methods. For example, the `SHAP` method (without the [CLS] and [SEP] tokens) has achieved 67.40% avg. F1-score with the heuristics, outperforming the vanilla `SHAP` method by *∼*10%. Similarly, the `SHAP` (with [CLS] and [SEP] tokens) achieved 68.73% F1-score with the heuristics, *∼*11% higher than the vanilla `SHAP` method. Finally, our best results have been achieved by using the vectors obtained from the `LIG` method for the 14^th^ hidden layer in the neural network model and applying the post-processing rules. This method resulted in the highest precision, highest recall and F1-score (overlap, exact matching, and average). For example, the 85.77% exact precision shows that in *∼*86% of the cases, detected spans are the actual triggers, and the 95.12% overlap precision reflects that in *∼*95% of the cases, detected spans overlap with the actual trigger spans, which is sufficient for the application point of view, because for large-scale execution, we prefer not to mark any spans if the method is not sure, but we want to make sure the detected spans overlap with the actual trigger spans. This method achieved 90.45% average precision, 67.43% average recall and 77.26% average F1-score, and it is used for large-scale trigger detection for STRING v12. These calculations penalize all methods in cases where multiple trigger spans are annotated for a relation, resulting to significantly lower recall. For that reason we manually calculated the overlap performance metrics of the best method (LIG+Heuristics) *without penalizing* for alternate trigger spans, to get an accurate picture of the performance on the development set and this resulted in a precision of 95.93%, a recall of 83.10% (increase by *∼*12%) and an F1-score of 89.06%.

#### Manual error analysis

Manual recalculation of the overlap performance metrics on the trigger test set yielded 92.80% precision, 79.93% recall and 85.88% F1-score. Detailed results are provided in Supplementary Material Section 6. A closer look in the results produced by the method shows that in cases where multiple words are valid as triggers, the method has a preference for punctuation (i.e. “-” or “/” in 35 cases) instead of whole words (e.g. “complex” in 5 cases). In cases where the method misdetects a trigger, by missing to predict the annotated trigger producing a FN, and by producing an incorrect trigger, i.e. a FP, there is no special pattern in the generated FPs, and the most frequent FN is the word “complex". FN `Complex_formation` relation predictions of the RE system, for which we had chosen not to run the trigger detection method as discussed above, inevitably result to FN predictions for the trigger detection method as well. Manual evaluation did not show any specific patterns pertaining these FNs. We also detected 4 annotation errors, where multiple triggers where valid and not all of them were annotated. This has been taken into consideration during the calculation of performance metrics above.

### Large-scale execution and integration in STRING v12

To perform relation extraction and trigger detection for STRING v12, we used all PubMed abstracts (as of August 2022) and all full-texts available in the PMC BioC text mining collection [Comeau et al., 2019] (as of April 2022). The entire corpus consists of 34,420,049 documents. We converted all documents into BRAT standoff format and obtained NER and `NEN Protein` annotations from Jensenlab tagger [Jensen, 2016] for both abstracts and full-text articles. 6,033,981 documents (3,604,037 abstracts and 2,429,944 full-texts) contained at least 2 protein NEs and were provided for `Complex_formation` relation prediction to the model with the best performance on the RE task. This resulted in predictions for over 1 billion NE pairs. From those 8,807,592 (less than 1%) are predicted as `Complex_formation` relationships. We then provide the *∼*8.8 million “positive” examples as input for the trigger detection pipeline. 7,127,119 of those examples have at least one trigger predicted. A tab-delimited file with scores produced by our RE model for all NE pairs, and the BRAT-formatted input and tab-delimited results from large-scale execution of the trigger detection system are provided through Zenodo.

Starting from version 11.5 of the STRING database [Szklarczyk et al., 2021], users gained access to a separate mode that provides a physical interaction subnetwork besides the broader functional association STRING network. Herein we have described a complete re-implementation of the text-mining pipeline that allows the detection of `Complex_formation` relationships (equivalent to “physical interactions” in STRING). The results from the large scale run are properly processed and incorporated in STRING. For details on post-processing please refer to the STRING publications [Szklarczyk et al., 2021, 2023]. In the physical interaction subnetwork of STRING v11.5, if there was evidence of a physical interaction between two unique Protein NEs based on text mining, the web interface enabled users to explore the publications supporting this interaction. This was accomplished by presenting actual text excerpts from biomedical articles, with the recognized `Protein` NEs highlighted. This feature served a dual purpose: users could personally evaluate the accuracy of automatically extracted relations, and they could also refer to the original articles for further in-depth reading.

Recognizing the potential for even greater utility, in the current version of STRING [Szklarczyk et al., 2023] it became desirable to have the system highlight the most significant word or phrase in the text snippet that explicitly or implicitly indicates the relation for each physical interaction extracted from that snippet. To achieve this, the results from the large-scale run of the trigger detection system were utilized. Supplementary Material Section 7 shows how text-mining results are presented for the physical interaction between two proteins in STRING v11.5 and v12.

## Conclusions

In this work, we present both ComplexTome, a corpus tailored for relation extraction of `Complex_formation` relationships among biomedical entities, and a relation extraction system that allows supervised training on the task, alongside large-scale execution on the biomedical literature. On top of relation extraction we also present a trigger detection method for `Complex_formation` relationships. Both the relation extraction and trigger detection methods achieve high levels of accuracy (F-score=82.8% and 85.9% on test set, respectively) and are available for use by the entire scientific community. We have meticulously manually evaluated both systems and integrated them into the latest version of the STRING database. As a result, we have not only augmented the database’s coverage of physical interactions, but also empowered users to explore and validate these relationships in the context of the original scientific articles. Overall, this project exemplifies the continuous evolution of text-mining capabilities in the era of language modeling and marks a significant milestone in enhancing our understanding of complex biological systems within the biomedical domain.

## Supporting information

Supplementary Section

## Acknowledgements

We thank the CSC – IT Center for Science for generous computational resources.

## Data availability

All resources introduced in this study are available under open licenses though Zenodo (https://doi.org/10.5281/zenodo.10693924) and GitHub (https://github.com/farmeh/ComplexTome_extraction).

