## Supplementary Section for "STRING-ing together protein complexes: corpus and methods for extracting physical protein interactions from the biomedical literature"

[†] Equal contribution

### 1. Implementation of marking and masking approach in the relation extraction system

Since a `Complex_formation` relation by definition is non-directional (i.e., `R(e1,e2)=R(e2,e1)`), an input document with $N$ NEs includes $N!/((N-2)!\times 2)$ candidate entity pairs, and for each pair, the system has to predict and assign a positive or a negative label. A typical document usually contains more than two entities (i.e., there are more than one pair in a typical document), therefore it is necessary to inform the classifier which two particular NEs constitute a pair at-a-time for label prediction. For this aim, we transform the text by encoding the entities in the input document, either using a marking approach or a masking approach. We use language model's "`unused`" tokens for this aim.

#### The marking approach:

In our marking approach, only the two focused entities (that constitute a pair at-a-time) are marked, but the texts of other entities in the text remain untouched. We use different "unused" tokens to mark the beginning and end of the focused entities, based on their type (e.g., `[unused1]` and `[unused2]` tokens are used to mark a `Protein` entity boundary, `[unused3]` and `[unused4]` are used to mark a `Chemical` boundary, etc). Therefore, our marking approach not only denotes which two entities constitute a pair for label detection, but it also denotes their type, providing maximum information to the classifier.

Following is an example sentence with three `Protein` entities and a `Protein_Family` entity and shows how the marking approach works. Note how different "unused" tokens are utilized to denote entity boundaries based on entity type.

Sentence:

"**GrpL**, a **Grb2-related adaptor protein**, interacts with **SLP-76** to regulate **nuclear factor of activated T cell** activation."

Entities:

`Protein` entities = {"**GrpL**", "**Grb2-related adaptor protein**", "**SLP-76**"}
`Protein_Family` entities = {"**nuclear factor of activated T cell**"}

There are six candidate pairs in the sentence which are transformed differently based on the focused entities:

- "[unused1]**GrpL**[unused2], a [unused1]**Grb2-related adaptor protein**[unused2], interacts with SLP-76 to regulate nuclear factor of activated T cell activation."

- "[unused1]**GrpL**[unused2], a Grb2-related adaptor protein, interacts with [unused1]**SLP-76**[unused2] to regulate nuclear factor of activated T cell activation."

- "[unused1]**GrpL**[unused2], a Grb2-related adaptor protein, interacts with SLP-76 to regulate [unused7]**nuclear factor of activated T cell**[unused8] activation."

- "GrpL, a [unused1]**Grb2-related adaptor protein**[unused2], interacts with [unused1]**SLP-76**[unused2] to regulate nuclear factor of activated T cell activation."

- "GrpL, a [unused1]**Grb2-related adaptor protein**[unused2], interacts with SLP-76 to regulate [unused7]**nuclear factor of activated T cell**[unused8] activation."

- "GrpL, a Grb2-related adaptor protein, interacts with [unused1]**SLP-76**[unused2] to regulate [unused7]**nuclear factor of activated T cell**[unused8] activation."

#### The masking approach

In the masking approach, the texts of the two focused entities in the text are always masked with `[unused1]` tokens, while **all other entities** in the text are masked with `[unused2]` tokens. Therefore, not only we hide the texts of all entities, but we also hide their types across the whole document. This is to ensure maximum generalization on unseen texts as the neural network model has to rely on and learn from non-entity words as context.

Following are the 6 transformations of the mentioned sentence, based on the masking approach:

- "**[unused1]**, a **[unused1]**, interacts with [unused2] to regulate [unused2] activation."
- "**[unused1]**, a [unused2], interacts with **[unused1]** to regulate [unused2] activation."
- "**[unused1]**, a [unused2], interacts with [unused2] to regulate **[unused1]** activation."

- "[unused2], a **[unused1]**, interacts with **[unused1]** to regulate [unused2] activation."
- "[unused2], a **[unused1]**, interacts with [unused2] to regulate **[unused1]** activation."
- "[unused2], a [unused2], interacts with **[unused1]** to regulate **[unused1]** activation."

#### A note on example generation:

For each candidate named-entity pair, after marking/masking the entities in the input text, we tokenize the transformed text into its corresponding sub-tokens (based on the vocabulary of a particular language model currently being used) and for each pair, we calculate their distance in sub-tokens (including the added markers in case of marking) and if they can fit into a window with a size smaller than or equal to the specified **MSL**, we generate a machine learning example (a sequence of tokens as a neural network input) for the pair.

We highlight that the same example generation method is used for positive/negative examples (i.e., during the training), and for unlabeled examples (i.e., during the prediction). Those longer examples not fitting into a window will be either discarded (in the case of training or predicting the unlabeled examples in large-scale prediction) or will be counted as False Negative (FN) predictions of the system (if there is a Complex_formation relation between their entities, in case of development/test set pairs).

### 2. List of transformer models and hyper-parameters used for building the relation extraction system

| | |
|---|---|
| Models | • BioBERT-base (only used for fast comparison of different training schemes) (download) (paper)<br>• BioBERT-large (download) (paper)<br>• RoBERTa-large-PM-M3-Voc (download) (paper) |
| Max sequence length | 128, 144, 160, 176, 192 |
| Learning rate | 2e-06, 3e-06, 4e-06, 5e-06 |
| Mini-batch size | • 5 (for the large models)<br>• 16 (for the bioBert-base model) |
| Number of epochs | 6,7,8,9,10,11,12 |

We used the BioBERT-base model for quick comparison of different training schemes (see here). For building the final system we tested BioBERT-large and RoBERTa-large-PM-M3-Voc models since these two models have recently achieved state-of-the-art on various tasks. Initial experiments showed that RoBERTa-large-PM-M3-Voc outperforms the BioBERT-large model, therefore, it was used for building the final system.

### 3. Different training schemes and initial evaluation results

The corpus contains **four** named-entity types (Protein, Chemical, Complex, Protein_Family/Family) and Complex_formation relationships can occur between any two entities mentioned in the text. For the real-world application for which the model was intended to be used (i.e., extracting Protein-Protein interactions for the STRING database v12), the system has to deal with texts including only Protein entities. Hence, to have a realistic optimization method for large-scale prediction, we filter-out all non-Protein entities and all Protein entities with the "blocklisted" attribute (and their relations) from the development and the test sets. In order to remove the Protein entities with the “blocklisted” attribute, we first convert them into a new entity type (Protein_BL) to allow for their easy removal when necessary.

We performed experiments with five different training schemes detailed below. Supplementary Table 1 shows an overview of each training scheme along with the highest f1-score achieved on the development set.

We used the BioBert-base model for all of the above training schemes (since it is a relatively small language model, thus requiring less computational resources, and we can get the results faster). For each training scheme, we run a full grid search to find the optimal values of hyper-parameters. For each hyper-parameter set, we repeat each experiment four times and calculate the average and standard deviation of the f1-score. Finally, we find and report the best obtained f1-score average and standard deviation. Following is the explanation of each approach.

**Training scheme 1-A:** In this approach, we use all available training data (2489 annotated Complex_formation relations among all entity types), and use the marking approach to denote the candidate entities in the input texts and their types. As Table1 shows, this approach yields the highest f1-score with an average of 81.10%and 0.2915 standard deviation.

**Training scheme 1-B:** This approach is similar to **1-A**, however, instead of marking the candidate entities, we mask all entity texts and their types, resulting in an average f1-score of 80.20%.

**Training scheme 2:** In this approach, we filter out all (i.e., delete) annotations for non-Protein entities and any annotated Complex_formation between them from the training set, and use the masking approach to hide all remaining entities and their types. This approach results in the minimum number of training examples

(1961), but is the most similar set to the reduced development set, i.e., the set that only contains `Protein` entities and interactions among them. This approach resulted in 79.28% average f1-score with 0.36 standard deviation. Please note that annotations for `Protein_BL`, `Complex`, `Chemical` and `Family` entities are deleted from the training .ann files, hence their texts are not masked in the experiment, and constitute part of the contexts if they appear in the window of `Protein-Protein` training examples.

**Training scheme 3:** In this approach, we only use entities of types `Protein`, `Complex` and `Protein_BL` (by converting the type of all `Complex` and `Protein_BL` entities to `Protein`), and then use the masking approach to mask the texts of all entities and their types (similar to Training scheme 1-B). As a result, `Family` and `Chemical` entities in context are retained, but no training examples for candidate pairs that either one or both of the entities are `Family` or `Chemical` are generated. The reasoning behind this training scheme is that in the scientific literature people tend to discuss `Complex` and `Protein_BL` entities in a similar manner as `Protein` entities, so there could be a benefit from using all of them while training, since this allows for a higher support for `Complex_formation` relationships (2117).

**Training scheme 4:** This approach is very similar to Training scheme 3, in the sense that only entities of types `Protein`, `Complex` and `Protein_BL` are masked and used to generate examples. The difference in this case is that annotations for `Family` and `Chemical` entities are completely removed and thus the text of these entities now constitutes part of the contexts if it appears within the window of a training example. This experiment was done to explore whether the text of `Family` and/or `Chemical` entities is important for the model towards predicting a `Complex_formation` relationship between two entities.

Out of all four schemes we have chosen to use **training scheme 1-B** for our relation extraction model training. There are two main reasons for this choice. Firstly, in comparison to training schemes 2,3 and 4, where masking is used as an encoding type, there is no statistically significant difference to the results obtained on the development set and the fact that we have the maximum amount of training data and maximum masking in this case, means that this is the most generalizable approach, out of these 4. When it comes to comparing with training scheme 1-A, the only difference is the approach used for encoding types between the two experiments. Again the difference is not statistically significant (i.e. the difference of the mean F-scores is within ±3std) and the masking approach is more generalizable to the open-world scenario, thus we have chosen training scheme 1-B for all subsequent experiments.

| Supplementary Table 1: Evaluation of different training schemes on the development set | | | | | | | | | |
|---|---|---|---|---|---|---|---|---|---|
| Exp# | Encoding type | Training pairs (Positive/Negatives) | Masked entities | Not masked entities | Removed annotations | Comments | Highest mean(F1) | std(F1) | Positive pairs |
| 1-A | Marking | Pair = (e1, e2) e1, e2 ∈ {Protein, Protein_BL, Complex, Chemical, Family} | | Protein, Protein_BL, Complex, Chemical, Family | | Maximum training data<br>No masking | 81.10 | 0.29 | 2489 |
| 1-B | Masking | Pair = (e1, e2) e1, e2 ∈ {Protein, Protein_BL, Complex, Chemical, Family} | Protein, Protein_BL, Complex, Chemical, Family | | | Maximum training data<br>Maximum masking | 80.20 | 0.60 | 2489 |
| 2 | Masking | Pair = (Protein, Protein) | Protein | | Protein_BL, Complex, Chemical, Family | Minimum training data<br>Minimum masking<br>similar to filtered dev set | 79.28 | 0.36 | 1961 |
| 3 | Masking | Pair = (e1, e2) e1, e2 ∈ {Protein, Protein_BL, Complex} | Protein, Protein_BL, Complex, Chemical, Family | | | More training data compared to Experiment 2<br>Maximum masking | 80.05 | 0.84 | 2117 |
| 4 | Masking | Pair=(e1, e2) e1, e2 ∈ {Protein, Protein_BL, Complex} | Protein, Protein_BL, Complex, | | Chemical, Family | More training data compared to Experiment 2<br>Less masking compared to Experiment 3 | 80.18 | 0.44 | 2117 |

### 4. Trigger Detection Methods

#### A. LIG-based trigger detection method

Our best relation extraction model was obtained by fine-tuning a pre-trained RoBERTa model (RoBERTa-large-PM-M3-Voc) on the relation extraction training set. We mainly focus on the outputs of the embedding layer and the outputs of the 24 hidden RoBERTa layers in this model. For each pair in the trigger development set, we feed the corresponding tokens as input to the model, and by applying the LIG method for all mentioned layers, we obtain 25 one-dimensional vectors, each representing the importance scores for the input tokens based on the outputs of a particular layer. By stacking the layers vertically, one can create a heatmap (rows denoting the layers, and columns denoting the tokens) and try to choose a layer (row in the heatmap) in which, the token with the highest score, is actually the trigger of the pair we are aiming to recognize.
The best layer might vary across different examples, thus we need a systematic approach for choosing the layer which yields the highest evaluation score when all predictions (i.e., predicted triggers for all positive pairs) are checked against the gold-standard (the annotated triggers in the trigger development set).

#### B. SHAP-based trigger detection method

Similarly to the LIG method, SHAP also yields a vector for the input tokens. For each pair, we simply feed the corresponding tokens as input to the model and apply the SHAP method, and then choose the token(s) with the highest score as the trigger(s). By repeating the process for all positive pairs, we obtain a prediction set which is then evaluated against the gold-standard. As an additional experiment, we try feeding the inputs ***with*** and ***without*** the [CLS] and [SEP] tokens, as initial experiments showed that the approaches produce slightly different results which demand further evaluation.

For both LIG and SHAP -based methods, we use the following approach: after obtaining a corresponding vector for a pair, we first discard the first and the last element of the vector (scores for the [CLS] and [SEP] tokens). Then we discard all “unused” tokens (which represent the entities) from the vector and then we choose the token(s) with the highest scores as the trigger(s) for that particular pair. By repeating the process for all positive pairs and for all layers, we obtain 25 LIG-based prediction sets (each based on choosing a particular layer as the best layer), and two SHAP-based prediction sets (with and without [CLS] and [SEP] tokens) and then we compare the prediction sets against the gold-standard and calculate evaluation scores.

#### C. Post-processing heuristic rules

Initial experiments showed that none of the two methods yield great evaluation scores, because sometimes tokens that are not actual triggers get the highest scores. To improve the results, we implemented the following post-processing heuristics. After a vector is calculated, and before selecting the token(s) with the highest score, we search all tokens in the vector and discard any token if one of the following rules applies:

- If the token is fully composed of white space(s) or punctuation mark(s) not found in the development set triggers ("-", ".","/" punctuation marks are actually valid triggers in the trigger development set).
- If \n or \t character is found inside the token text.
- If the token is a "." character and it is the first or the last token in the sequence (after removing [CLS] and [SEP] tokens)
- If the token is "." character, is located in the middle of the snippet, and it is not in the "[unused1].[unused1]" pattern.
- If the token belongs to a list of closed class words, such as pronouns, prepositions, conjugations, etc. The complete list of closed class words used is given in Supplementary Material Section 4E.

The aforementioned rules are obtained by inspecting the heatmaps and SHAP outputs for the (positive) pairs in the trigger development set. Finally, it is worth mentioning that our method can provide multiple disjoint trigger spans for a `Complex_formation` relationship, since we are finding the token(s) in a vector with the maximum score (i.e., if there are multiple tokens with the same highest score, they will all be returned as recognized triggers).

#### D. An example output for LIG and SHAP methods

The following shows the outputs of SHAP and part of the heatmap obtained by running layer integrated gradients with Captum. Please note that the two candidate named-entities are replaced with [unused1] tokens.

**SHAP output:**

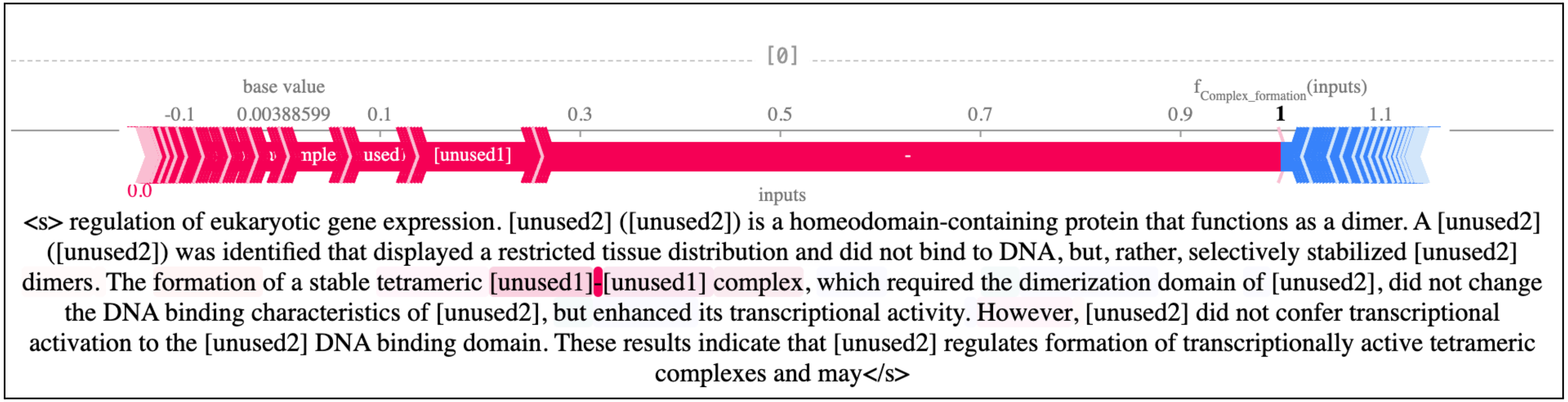


As SHAP result shows, the highest scores belong to ‘-’, [unused1] and ‘complex’ tokens. This is logical, as removing [unused1] tokens from the input should change the prediction outcome, i.e., the positive label (Complex_formation). This is the reason why we remove [unused1] tokens from the output of SHAP and LIG methods before finding the token with the highest score.

The following is a part of the heatmap, obtained by running the LIG algorithm on the same sentence. As we notice, there are a lot of high hits around the [unused1] tokens, as well as the ‘-’ token and ‘Complex’ token. As we notice, lower layers in the network (closer to the input), have higher numbers for the ‘-’ and ‘complex’ tokens, but higher layers (closer to the output), have higher scores for the [unused1] tokens. Also note that the numbers for each row do **not** sum up to 100.

| | ĠThe | Ġformation | Ġof | Ġa | Ġstable | Ġtetrameric | Ġ | [unused1] | - | [unused1] | Ġcomplex | , | Ġwhich | Ġrequired | Ġthe | Ġdimerization | Ġdomain |
|---|---|---|---|---|---|---|---|---|---|---|---|---|---|---|---|---|---|
| 23 | 0.000 | 0.000 | 0.000 | 0.000 | 0.000 | 0.000 | 0.000 | 0.000 | 0.000 | 0.000 | 0.000 | 0.000 | 0.000 | 0.000 | 0.000 | 0.000 | 0.000 |
| 22 | 1.531 | 3.358 | 4.568 | 6.362 | 2.490 | 5.186 | 9.769 | 9.920 | 9.712 | 10.513 | 7.224 | 3.558 | 0.717 | 0.392 | 0.453 | 0.432 | 0.357 |
| 21 | 0.166 | 2.214 | 3.321 | 8.529 | 1.309 | 6.305 | 31.516 | 34.723 | 30.925 | 39.330 | 12.245 | 0.316 | 0.337 | 0.307 | 0.248 | 0.253 | 0.290 |
| 20 | -0.600 | 1.099 | 1.568 | 5.060 | 0.626 | 1.849 | 25.754 | 39.092 | 50.523 | 69.007 | 7.805 | -3.590 | 0.050 | 0.077 | -0.109 | -0.139 | 0.020 |
| 19 | -1.008 | 0.586 | 0.588 | 1.735 | 0.354 | 1.537 | 19.917 | 36.754 | 46.165 | 77.417 | 7.517 | -7.077 | 0.158 | 0.214 | 0.109 | 0.164 | 0.437 |
| 18 | -0.524 | -0.441 | -1.075 | -1.297 | -0.280 | -0.317 | 9.072 | 58.563 | 27.092 | 75.014 | -0.713 | -3.248 | -0.170 | 0.086 | 0.310 | 0.155 | 0.159 |
| 17 | -0.422 | -0.426 | -0.451 | -0.341 | -0.667 | -1.011 | 2.003 | 74.995 | 8.566 | 65.026 | -0.847 | -2.921 | -0.146 | 0.017 | 0.161 | -0.123 | -0.090 |
| 16 | -0.732 | 2.142 | 1.444 | 0.787 | 0.078 | -2.099 | 4.496 | 73.162 | 29.060 | 59.009 | 2.967 | -1.683 | 0.104 | -0.451 | -0.003 | 0.040 | -0.058 |
| 15 | -1.702 | 1.460 | 2.146 | 2.287 | 2.549 | 2.247 | 3.443 | 59.614 | 60.360 | 45.958 | 21.414 | -1.110 | -0.469 | -0.144 | -0.173 | 0.095 | -0.116 |
| 14 | -2.352 | 1.425 | 1.776 | 2.583 | 4.649 | 7.468 | 7.370 | 55.175 | 45.217 | 58.128 | 32.410 | -1.677 | -1.243 | -1.132 | -0.490 | 0.127 | -0.339 |
| 13 | -3.934 | -0.383 | 0.963 | 3.722 | 4.673 | 7.945 | 10.543 | 35.450 | 83.759 | 25.693 | 26.498 | -2.475 | -1.211 | 0.328 | -0.527 | 0.006 | -0.497 |
| 12 | -4.516 | 0.183 | 1.168 | 4.041 | 7.488 | 11.607 | 17.723 | 24.354 | 63.085 | -1.438 | 52.631 | -3.128 | -1.698 | -2.574 | -1.188 | 0.504 | -0.522 |
| 11 | -1.618 | 0.186 | -0.054 | 1.196 | 2.787 | 8.131 | 12.668 | 41.058 | 76.323 | 32.737 | 16.954 | -0.376 | -3.265 | 0.575 | 0.160 | 0.121 | 0.091 |
| 10 | 4.031 | 2.401 | 3.598 | 6.297 | 7.255 | 10.582 | 5.686 | -29.344 | 43.293 | -6.746 | 27.116 | -0.452 | 0.066 | 0.433 | 0.205 | -0.453 | -0.688 |
| 9 | 1.515 | 2.374 | 3.740 | 4.440 | 8.103 | 12.772 | 12.448 | 14.424 | 45.284 | 16.353 | 58.690 | 5.942 | 2.550 | -0.851 | 0.088 | -2.133 | -1.190 |
| 8 | 0.693 | 2.635 | 1.586 | 1.492 | 7.467 | 3.035 | 15.279 | -23.029 | 65.835 | -40.806 | 42.841 | 10.222 | 3.888 | 3.987 | 0.170 | -2.146 | -2.589 |
| 7 | 0.778 | 2.096 | 1.154 | 2.103 | 6.557 | 3.309 | 9.712 | -16.883 | 51.660 | -25.789 | 46.193 | 3.134 | 2.661 | 6.421 | 0.591 | -1.317 | -4.095 |
| 6 | 0.866 | -1.625 | 1.156 | 0.517 | 6.418 | 2.126 | 6.597 | -33.233 | 38.380 | 28.862 | 29.817 | 0.475 | -0.008 | 1.491 | -0.498 | -2.102 | -0.315 |
| 5 | 3.104 | 1.721 | 2.600 | 0.827 | 3.370 | -7.618 | 11.408 | -33.038 | 49.387 | -1.760 | 56.628 | 7.129 | -2.601 | 2.540 | 0.537 | -3.642 | -3.424 |
| 4 | 1.023 | -4.176 | -2.734 | -0.457 | 10.280 | -10.089 | -7.889 | -37.395 | 44.649 | 2.696 | 65.498 | 15.834 | 1.335 | 7.550 | 0.307 | -3.733 | -0.545 |
| 3 | -3.857 | -0.795 | -1.407 | 1.506 | 7.086 | -1.291 | -8.114 | -50.828 | 65.311 | -18.844 | 39.592 | -3.819 | 2.623 | -2.522 | -0.387 | -4.126 | -0.715 |
| 2 | -5.860 | -1.289 | -4.252 | 0.215 | 1.370 | 2.451 | -0.682 | -36.576 | 32.338 | -20.239 | 10.954 | -1.307 | 0.409 | 1.522 | -4.939 | -0.090 | -1.412 |
| 1 | -5.761 | 5.488 | -4.735 | 12.372 | 7.468 | 15.919 | -35.058 | -32.376 | 47.344 | -8.836 | 26.688 | -12.632 | -33.351 | 23.934 | 5.317 | -3.435 | -3.390 |
| 0 | 1.458 | -2.533 | -2.940 | -1.930 | 6.293 | 9.087 | -10.644 | 14.931 | 60.572 | 10.519 | 41.828 | -0.500 | 0.248 | -0.014 | 2.446 | -0.614 | 0.794 |
| embeddings | -2.905 | 0.707 | 4.800 | 5.773 | 5.332 | -1.487 | 0.904 | -0.386 | 7.242 | 6.091 | 11.238 | 16.226 | -1.650 | 1.032 | -2.435 | 0.429 | 0.244 |

### E. Closed class words in trigger word detection

```
CCWords = {'&', "'cause", "'n", "'n'", "'til", 'I', 'a', 'aboard', 'about',
'above', 'across', 'after', 'against', 'ago','albeit', 'all', 'along',
'alongside', 'although', 'always', 'am', 'amid', 'among', 'amongst', 'an',
'and', 'any', 'anybody', 'anyhow', 'anyone', 'anything', 'anytime',
'anyway', 'anywhere', 'are', 'around', 'as', 'astride', 'at','atop', 'be',
'because', 'been', 'before', 'behind', 'being', 'below', 'beneath',
'beside', 'besides', 'between','beyond', 'billion', 'billionth', 'both',
'but', 'by', 'can', 'cannot', 'could', 'de', 'despite', 'did', 'do','does',
'doing', 'done', 'down', 'during', 'each', 'eight', 'eighteen',
'eighteenth', 'eighth', 'eightieth', 'eighty','either', 'eleven',
'eleventh', 'en', 'enough', 'et', 'every', 'everybody', 'everyone',
'everything', 'everywhere','except', 'few', 'fewer', 'fifteen',
'fifteenth', 'fifth', 'fiftieth', 'fifty', 'first', 'five', 'for',
'fortieth','forty', 'four', 'fourteen', 'fourteenth', 'fourth', 'from',
'had', 'has', 'have', 'having', 'he', 'her', 'here', 'hers', 'herself',
'him', 'himself', 'his', 'how', 'hundred', 'hundredth', 'if', 'in',
'inside', 'into', 'is','it', 'its', 'itself', 'least', 'less', 'lest',
'like', 'little', 'many', 'may', 'me', 'might', 'million', 'millionth',
'mine', 'minus', 'more', 'most', 'much', 'must', 'my', 'myself', 'near',
'neither', 'never', 'next', 'nine', 'nineteen', 'nineteenth', 'ninetieth',
'ninety', 'ninth', 'no', 'nobody', 'none', 'nor', 'not', 'nothing',
'notwithstanding', 'now', 'nowhere', 'of', 'off', 'on', 'one', 'oneself',
'onto', 'opposite', 'or', 'our', 'ours', 'ourselves', 'out', 'outside',
'over', 'par', 'past', 'per', 'plus', 'post', 'second', 'seven',
'seventeen', 'seventeenth', 'seventh', 'seventieth', 'seventy', 'shall',
'she', 'should', 'since', 'six', 'sixteen', 'sixteenth', 'sixth',
'sixtieth', 'sixty', 'so', 'some', 'somebody', 'somehow', 'someone',
'something', 'sometime', 'somewhere', 'ten', 'tenth', 'than', 'that',
'the', 'their', 'theirs', 'them', 'themselves', 'then', 'there', 'these',
'they', 'third', 'thirteen', 'thirteenth', 'thirtieth', 'thirty', 'this',
'those', 'though', 'thousand', 'thousandth', 'three', 'through',
'throughout', 'till', 'times', 'to', 'too', 'toward', 'towards', 'twelfth',
'twelve', 'twentieth', 'twenty', 'two', 'under', 'underneath', 'unless',
'unlike', 'until', 'unto', 'up', 'upon', 'us', 'v.', 'versus', 'via',
'vs.', 'was', 'we', 'were', 'what', 'when', 'where', 'whereas', 'whether',
'which', 'while', 'who', 'whom', 'whose', 'why', 'will', 'willing', 'with',
'within', 'without', 'worth', 'would', 'yes', 'yet', 'you', 'your',
'yours', 'yourself', 'yourselves', 'zero'}
```

### 5. Table with results of relation extraction error analysis results for the best model on the test set

*FN: False Negative, FP: False Positive*

| PubMed ID | FP/FN | Error Type |
|---|---|---|
| 14562105 | FP | co-reference resolution |
| 16177062 | FP | co-reference resolution |
| 16177062 | FP | co-reference resolution |
| 16209941 | FP | co-reference resolution |
| 16209941 | FP | co-reference resolution |
| 16209941 | FP | co-reference resolution |
| 16209941 | FP | co-reference resolution |
| 16260776 | FP | co-reference resolution |
| 16260776 | FP | co-reference resolution |
| 20406818 | FP | co-reference resolution |
| 23022657_23 | FP | co-reference resolution |
| 10446169 | FN | co-reference resolution |
| 15485920 | FN | co-reference resolution |
| 15485920 | FN | co-reference resolution |
| 16177062 | FN | co-reference resolution |
| 16908542 | FN | co-reference resolution |
| 16908542 | FN | co-reference resolution |
| 16908542 | FN | co-reference resolution |
| 20406818 | FN | co-reference resolution |
| 23022657_23 | FN | co-reference resolution |

| PubMed ID | FP/FN | Error Type |
|---|---|---|
| 15449939 | FP | convoluted text excerpt |
| 1719979 | FP | convoluted text excerpt |
| 1719979 | FP | convoluted text excerpt |
| 17435760 | FP | convoluted text excerpt |
| 18317453 | FP | convoluted text excerpt |
| 24498436 | FP | convoluted text excerpt |
| 25569479_19 | FP | convoluted text excerpt |
| 9858532 | FP | convoluted text excerpt |
| 11520069 | FN | convoluted text excerpt |
| 11520069 | FN | convoluted text excerpt |
| 11520069 | FN | convoluted text excerpt |
| 11520069 | FN | convoluted text excerpt |
| 11520069 | FN | convoluted text excerpt |
| 11520069 | FN | convoluted text excerpt |
| 11867544 | FN | convoluted text excerpt |
| 12151385 | FN | convoluted text excerpt |
| 12151385 | FN | convoluted text excerpt |
| 14670962 | FN | convoluted text excerpt |
| 15449939 | FN | convoluted text excerpt |
| 16209941 | FN | convoluted text excerpt |

| | | |
|---|---|---|
| 16908542 | FP | ambiguous keyword |
| 19276361 | FP | ambiguous keyword |
| 20797779_6 | FP | ambiguous keyword |
| 22958824_7 | FP | ambiguous keyword |
| 28515143_17 | FP | ambiguous keyword |
| 9195882 | FP | ambiguous keyword |
| 9367446 | FP | ambiguous keyword |
| 9367446 | FP | ambiguous keyword |
| 10433269 | FN | ambiguous keyword |
| 21236256 | FN | ambiguous keyword |
| 21554248 | FN | ambiguous keyword |
| 21554248 | FN | ambiguous keyword |
| 22086907_3 | FN | ambiguous keyword |
| 22355353 | FN | ambiguous keyword |
| 23499533_3 | FN | ambiguous keyword |
| 25602519 | FN | ambiguous keyword |
| 27751725 | FN | ambiguous keyword |
| 27751725 | FN | ambiguous keyword |
| 27751725 | FN | ambiguous keyword |
| 27751725 | FN | ambiguous keyword |
| 28515143_17 | FN | ambiguous keyword |
| 9115214 | FN | ambiguous keyword |
| 9115214 | FN | ambiguous keyword |
| 9195882 | FN | ambiguous keyword |
| 9348293_33 | FN | ambiguous keyword |

| | | |
|---|---|---|
| 16209941 | FN | convoluted text excerpt |
| 19228687 | FN | convoluted text excerpt |
| 19228687 | FN | convoluted text excerpt |
| 21236256 | FN | convoluted text excerpt |
| 21807881 | FN | convoluted text excerpt |
| 21827752 | FN | convoluted text excerpt |
| 9348293_33 | FN | convoluted text excerpt |
| 9348293_33 | FN | convoluted text excerpt |
| 9348293_33 | FN | convoluted text excerpt |
| 9195882 | FP | convoluted text excerpt |
| 10770935 | FN | convoluted text excerpt |
| 21347367 | FN | convoluted text excerpt |
| 7524088 | FN | convoluted text excerpt |
| 8505295 | FN | convoluted text excerpt |
| 8505295 | FN | convoluted text excerpt |
| 10585430 | FP | convoluted text excerpt |
| 11416152 | FP | convoluted text excerpt |
| 11416152 | FP | convoluted text excerpt |
| 11416152 | FP | convoluted text excerpt |
| 11416152 | FP | convoluted text excerpt |
| 12135708 | FP | convoluted text excerpt |
| 10446169 | FN | convoluted text excerpt |
| 11416152 | FN | convoluted text excerpt |
| 11416152 | FN | convoluted text excerpt |
| 11416152 | FN | convoluted text excerpt |

| | | |
|---|---|---|
| 9506992 | FN | ambiguous keyword |
| 26651479_12 | FP | ambiguous keyword |
| 29283431 | FP | ambiguous keyword |
| 8407894 | FP | ambiguous keyword |
| 8407894 | FP | ambiguous keyword |
| 8407894 | FP | ambiguous keyword |
| 8407894 | FP | ambiguous keyword |
| 8505295 | FP | ambiguous keyword |
| 8505295 | FP | ambiguous keyword |
| 8505295 | FP | ambiguous keyword |
| 8505295 | FP | ambiguous keyword |
| 29666278 | FN | ambiguous keyword |
| 8407894 | FN | ambiguous keyword |
| 8407894 | FN | ambiguous keyword |
| 8407894 | FN | ambiguous keyword |
| 8407894 | FN | ambiguous keyword |
| 8407894 | FN | ambiguous keyword |
| 8407894 | FN | ambiguous keyword |
| 8407894 | FN | ambiguous keyword |
| 11416152 | FP | annotation error |
| 11416152 | FP | annotation error |
| 12151385 | FP | annotation error |
| 16908542 | FP | annotation error |
| 17194709 | FP | annotation error |
| 17194709 | FP | annotation error |
| 20463880 | FP | annotation error |

| | | |
|---|---|---|
| 11521196 | FN | convoluted text excerpt |
| 11521196 | FN | convoluted text excerpt |
| 12151385 | FN | convoluted text excerpt |
| 17572495 | FN | convoluted text excerpt |
| 21554248 | FN | convoluted text excerpt |
| 21554248 | FN | convoluted text excerpt |
| 23533635 | FN | convoluted text excerpt |
| 23533635 | FN | convoluted text excerpt |
| 8521815 | FN | convoluted text excerpt |
| 14562105 | FP | rare keyword |
| 8505295 | FP | rare keyword |
| 10666337 | FN | rare keyword |
| 11520069 | FN | rare keyword |
| 11520069 | FN | rare keyword |
| 11713274 | FN | rare keyword |
| 11897782 | FN | rare keyword |
| 15223 | FN | rare keyword |
| 16209941 | FN | rare keyword |
| 16717095 | FN | rare keyword |
| 16717095 | FN | rare keyword |
| 16908542 | FN | rare keyword |
| 16908542 | FN | rare keyword |
| 16912044 | FN | rare keyword |
| 17194709 | FN | rare keyword |
| 17194709 | FN | rare keyword |
| 18628300 | FN | rare keyword |

| | | |
|---|---|---|
| 20463880 | FP | annotation error |
| 20463880 | FP | annotation error |
| 24498436 | FP | annotation error |
| 25938661_25 | FP | annotation error |
| 11874917 | FN | annotation error |
| 11897782 | FN | annotation error |
| 11897782 | FN | annotation error |
| 11897782 | FN | annotation error |
| 16738327 | FN | annotation error |
| 16912044 | FN | annotation error |
| 16912044 | FN | annotation error |
| 16912044 | FN | annotation error |

| | | |
|---|---|---|
| 19276361 | FN | rare keyword |
| 2002555 | FN | rare keyword |
| 20129058 | FN | rare keyword |
| 20129058 | FN | rare keyword |
| 20129058 | FN | rare keyword |
| 23533635 | FN | rare keyword |
| 25602519 | FN | rare keyword |
| 27120157 | FN | rare keyword |

### 6. Table with results of trigger word detection error analysis results for the best model on the test set

The first part of the results is the PMID of the document (e.g. 10080948) followed by the two entities (e.g. T13 & T14 for the 2nd cell in the first column) for which the trigger words annotations have been made in the provided text snippet. *4 examples (Table B) are counted as TP due to acceptable alternative annotations detected by the annotator and thus only 17 are counted as FP+FN
*FN: False Negative, FP: False Positive*

A) Correct trigger word detected in documents with multiple triggers (TP) (**Total count: 63)**

| PMID_EID1_EID2 | detected | not detected |
|---|---|---|
| 10080948_T13_T14 | - | interaction |
| 10096561_T28_T29 | - | interactions |
| 10229231_T11_T12 | / | interaction |
| 10229231_T3_T4 | / | interaction |
| 10229231_T3_T5 | / | interaction |
| 10395652_T18_T8 | complex | recruited |
| 10395652_T19_T8 | complex | recruited |
| 10395652_T5_T6 | associates | complex |
| 10704443_5_T2_T3 | complex | comprises |
| 10704443_5_T2_T4 | complex | comprises |
| 10704443_5_T2_T5 | complex | comprises |
| 10704443_5_T3_T4 | complex | comprises |
| 10704443_5_T3_T5 | complex | comprises |
| 10704443_5_T4_T5 | complex | comprises |
| 12024017_T13_T2 | / | complex |
| 12024017_T21_T10 | / | complex |
| 12897057_T11_T12 | complex | / |
| 12897057_T14_T15 | / | complex |
| 12897057_T18_T19 | / | complex |
| 15225555_T12_T13 | targets | yeast two-hybrid |
| 15225555_T12_T14 | targets | yeast two-hybrid |

| | | |
|---|---|---|
| 1700011_T5_T6 | / | heterodimers |
| 1700011_T7_T8 | / | heterodimers |
| 1763325_T6_T7 | - | complex |
| 19141288_18_T1_T2 | complex | / |
| 19141288_18_T7_T8 | complex | / |
| 19141288_18_T20_T21 | saturation | Kd |
| 19656901_T38_T8 | - | associations |
| 19656901_T7_T37 | - | associations |
| 20541251_2_T7_T12 | interaction | - |
| 21685921_32_T23_T1 | - | complexes |
| 21685921_32_T24_T3 | - | complexes |
| 22210188_T14_T5 | partner | yeast two-hybrid |
| 22210188_T14_T6 | partner | yeast two-hybrid |
| 22210188_T4_T5 | partner | yeast two-hybrid |
| 22210188_T4_T6 | partner | yeast two-hybrid |
| 22210188_T7_T8 | interaction | immunoprecipitation |
| 22675546_10_T11_T2 | coimmunoprecipitated | complex |
| 22675546_10_T14_T4 | complex | - |
| 22675546_10_T1_T2 | coimmunoprecipitated | complex |
| 22675546_10_T9_T2 | coimmunoprecipitated | complex |
| 24465968_22_T8_T9 | - | complex |
| 24498054_T19_T1 | / | complex |
| 24498054_T22_T6 | / | complex |
| 26996158_T15_T16 | - | complex |
| 27044741_T33_T25 | - | complex |
| 7492771_T11_T12 | / | heterodimers |
| 7492771_T15_T16 | / | heterodimers |
| 7492771_T5_T6 | / | heterodimers |
| 7537762_T10_T11 | binding | ligand |
| 8098618_T18_T19 | - | heterodimer |
| 8098618_T21_T22 | - | heterodimer |
| 8626752_T1_T2 | association | complexes |
| 8626752_T1_T3 | association | complexes |
| 8626752_T6_T7 | - | heterocomplexes |
| 9512491_T17_T18 | - | complex |

| | | |
|---|---|---|
| 9512491_T17_T19 | - | -<br>complex |
| 9512491_T18_T19 | - | complex |
| 9512491_T7_T17 | - | complex |
| 9512491_T7_T18 | - | -<br>complex |
| 9512491_T7_T19 | - | -<br>complex |
| 9733846_T11_T12 | - | interaction |
| 9733846_T15_T16 | - | interaction |

B) Word detected not a trigger (FP+FN) (**Total count: 21)**

| PMID_EID1_EID2 | detected | not detected |
|---|---|---|
| 12093729_T4_T41 | MP | in<br>complex |
| 12093729_T6_T41 | MP | in<br>complex |
| 1700011_T3_T4 | together | heterodimeric |
| 17452446_T4_T16 | forms | complex |
| 17452446_T4_T17 | forms | complex |
| 17452446_T4_T5 | forms | complex |
| 17452446_T4_T6 | forms | complex |
| 17452446_T8_T9 | components | complex |
| 19150429_T1_T2 | - | interacting |
| 19217404_T18_T19 | form | hexameric |
| 19217404_T18_T21 | form | hexameric |
| 19217404_T19_T21 | form | hexameric |
| 20541251_2_T10_T6* | target | bind<br>complex |
| 20541251_2_T17_T1 | binding | recruitment |
| 20541251_2_T8_T6* | target | bind<br>complex |
| 22904036_T10_T13 | including | interacted |
| 22904036_T11_T13 | including | interacted |
| 22904036_T3_T4 | including | interacting partners |
| 22904036_T3_T5 | including | interacting partners |
| 8626752_T12_T13* | - | complex |
| 9751051_T1_T2* | complex | bound |

C) No trigger word detected in documents with one trigger word (FN) (**Total count: 34)**

| PMID_EID1_EID2 | not detected |
|---|---|
| 15314162_T9_T10 | binding |
| 17452446_T11_T1 | interaction |
| 17452446_T11_T12 | interaction |
| 17452446_T12_T1 | interaction |
| 19150429_T6_T8 | interacts |
| 19217404_T22_T26 | receive |
| 19414608_38_T7_T8 | interaction |
| 20541251_2_T11_T15 | recruitment |
| 21063388_2_T7_T9 | interacting |
| 21063388_2_T9_T10 | interacting |
| 21907821_T10_T38 | binding |
| 21907821_T10_T39 | binding |
| 22056990_T10_T3 | target |
| 22160715_T8_T4 | binding |
| 22160715_T8_T5 | binding |
| 22160715_T8_T6 | binding |
| 22579246_3_T1_T13 | interact |
| 22579246_3_T1_T3 | interact |
| 22579246_3_T2_T13 | interact |
| 22579246_3_T2_T3 | interact |
| 26675234_T12_T14 | binds |
| 27044741_T23_T24 | - |
| 27044741_T28_T8 | - |
| 27044741_T30_T14 | - |
| 27044741_T5_T6 | - |
| 27044741_T7_T4 | - |
| 8649779_T10_T11 | subunits |
| 8649779_T10_T12 | subunits |
| 8649779_T10_T13 | physically interact |
| 8649779_T11_T12 | subunits |
| 9171108_T3_T4 | associated |
| 9356494_T9_T11 | associated |

| | |
|---|---|
| 9512491_T27_T28 | effector |
| 9990065_T10_T11 | radioligand binding |

D) No trigger word detected in documents with multiple trigger words (FN) **Total count: 4**

| PMID_EID1_EID2 | not detected |
|---|---|
| 14707132_T28_T30 | two-hybrid screen<br>target |
| 14707132_T29_T30 | two-hybrid screen<br>target |
| 22675546_10_T4_T5 | recruitment<br>complex |
| 27462423_T11_T12 | /<br>complex |

Correct trigger word detected in documents with one trigger word (TP) (**Total count: 152)**

10080948_T10_T11, 10080948_T1_T2, 10080948_T3_T4, 10080948_T6_T7, 10080948_T8_T9, 10395652_T9_T20, 11959841_T18_T4, 12093729_T10_T11, 12093729_T10_T44, 12093729_T17_T45, 12223469_T14_T15, 12223469_T16_T17, 12223469_T16_T18, 12223469_T1_T2, 12223469_T24_T25, 12223469_T30_T31, 12223469_T6_T7, 12223469_T9_T10, 14973287_T17_T20, 14973287_T17_T5, 14973287_T17_T6, 15107829_T3_T4, 15107829_T3_T5, 15107829_T8_T9, 15150273_T23_T24, 15150273_T29_T2, 15150273_T37_T11, 15150273_T41_T16, 15314162_T8_T9, 16546083_T19_T20, 16546083_T5_T6, 16787403_T19_T5, 16787403_T6_T7, 17452446_T14_T4, 17452446_T15_T4, 17452446_T5_T6, 17666399_T13_T30, 17666399_T7_T22, 17666399_T7_T23, 18029035_T14_T3, 18029035_T4_T6, 18029035_T4_T7, 18455122_T5_T6, 18455122_T5_T7, 19141288_18_T14_T15, 19141288_18_T3_T4, 19150429_T11_T2, 19150429_T11_T3, 19150429_T1_T3, 19150429_T6_T7, 19217404_T11_T12, 19217404_T11_T14, 19217404_T22_T23, 19217404_T26_T28, 19414608_38_T7_T9, 19414608_38_T8_T9, 20472562_T12_T15, 20472562_T12_T16, 20472562_T13_T15, 20472562_T13_T16, 20472562_T18_T20, 20472562_T19_T20, 20541251_2_T11_T22, 20541251_2_T17_T18, 20541251_2_T24_T9, 21063388_2_T12_T13, 21063388_2_T15_T16, 21063388_2_T19_T20, 21063388_2_T3_T5, 21063388_2_T3_T6, 21063388_2_T7_T8, 21985068_5_T2_T17, 21985068_5_T2_T3, 21985068_5_T2_T4, 21985068_5_T2_T5, 22009753_T9_T10, 22160715_T16_T1, 22210188_T11_T12, 22210188_T1_T13, 22210188_T1_T3, 22210188_T2_T13, 22210188_T2_T3, 22210188_T9_T10, 22675546_10_T17_T7, 22675546_10_T18_T7, 22675546_10_T3_T12, 22675546_10_T3_T13, 22675546_10_T6_T15, 22675546_10_T8_T20, 22842725_T3_T16, 22842725_T3_T4, 22904036_T10_T12, 22904036_T10_T14, 22904036_T11_T12, 22904036_T11_T14, 22904036_T9_T10, 22904036_T9_T11, 23446637_T1_T2, 23446637_T8_T16, 23446637_T8_T9, 23498974_T17_T18, 23498974_T17_T19, 24498054_T20_T2, 26675234_T12_T13, 26972597_T27_T28, 26972597_T32_T35, 26972597_T32_T36, 26996158_T10_T11, 26996158_T14_T15, 27334688_T6_T7, 27462423_T10_T11, 27462423_T10_T12, 27462423_T9_T18, 7492771_T1_T2, 7747417_T7_T8, 7747417_T9_T10, 8108127_T20_T21, 8108127_T20_T22, 8156587_T10_T5, 8626752_T10_T11, 8626752_T14_T15, 8626752_T14_T16, 8626752_T17_T19, 8626752_T18_T19, 8626752_T8_T9, 8649779_T11_T13, 8649779_T12_T13, 9171108_T11_T12, 9171108_T13_T14, 9171108_T1_T4, 9171108_T2_T4, 9171108_T9_T10, 9233773_T14_T12, 9233773_T16_T5, 9233773_T16_T6, 9353251_T12_T13, 9353251_T8_T9, 9356494_T12_T13, 9356494_T1_T2, 9356494_T9_T10, 9512491_T25_T26, 9751051_T3_T4, 9794375_T16_T17, 9794375_T16_T18, 17452446_T16_T5, 17452446_T16_T6, 17452446_T17_T5, 17452446_T17_T6, 19141288_18_T18_T19, 26996158_T14_T16, 9233773_T14_T13, 9233773_T14_T4

### 7. Text-mining results for the physical interaction between two proteins in STRING v11.5 and v12

Example of text excerpts supporting a text-mined physical protein interaction between TrpA and TrpB from the web interfaces of STRING v11.5 to v12. The same publication is shown for better comparison between the two versions. Trigger words are highlighted in v12.

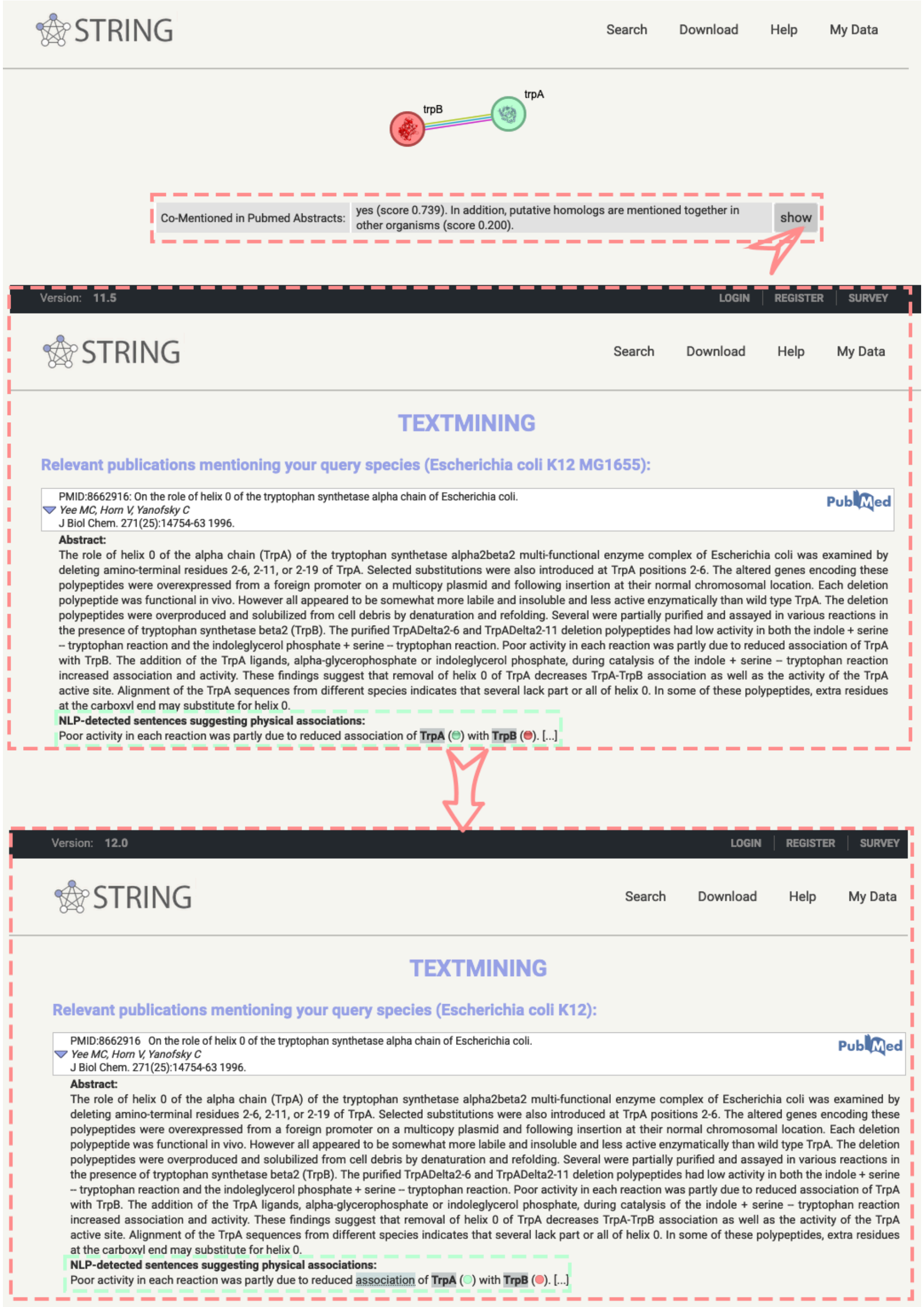
